# A Structure-based B-cell Epitope Prediction Model Through Combing Local and Global Features

**DOI:** 10.1101/2021.07.13.452188

**Authors:** Shuai Lu, Yuguang Li, Qiang Ma, Xiaofei Nan, Shoutao Zhang

**Affiliations:** School of Information Engineering, Zhengzhou University, Zhengzhou, China; College of Economics and Management, Zhengzhou University of Light Industry, Zhengzhou, China; School of Life Science, Zhengzhou University, Zhengzhou, China

**Keywords:** Attention, Bi-LSTM, B-cell epitope prediction, GCN, SARS-Cov-2, structure-based

## Abstract

B-cell epitopes (BCEs) are a set of specific sites on the surface of an antigen that binds to an antibody produced by B-cell. The recognition of BCEs is a major challenge for drug design and vaccines development. Compared with experimental methods, computational approaches have strong potential for BCEs prediction at much lower cost. Moreover, most of the currently methods focus on using local information around target residue without taking the global information of the whole antigen sequence into consideration. We propose a novel deep leaning method through combing local features and global features for BCEs prediction. In our model, two parallel modules are built to extract local and global features from the antigen separately. For local features, we use Graph Convolutional Networks(GCNs) to capture information of spatial neighbors of a target residue. For global features, Attention-Based Bidirectional Long Short-Term Memory(Att-BLSTM) networks are applied to extract information from the whole antigen sequence. Then the local and global features are combined to predict BCEs. The experiments show that the proposed method achieves superior performance over the state-of-the-art BCEs prediction methods on benchmark datasets. Also, we compare the performance differences between data with or without global features. The experimental results show that global features play an important role in BCEs prediction. Our detailed case study on the BCEs prediction for SARS-Cov-2 receptor binding domain confirms that our method is effective for predicting and clustering true BCEs.

## 1 INTRODUCTION

The humoral immune system protects the body from foreign objects like bacteria and viruses by developing B-cells and producing antibodies (1). Antibodies play a crucial role in immune response through recognizing and binding the disease-causing agents, called antigen. B-cell epitopes (BCEs) are a certain region on the antigen surface that are bound by an antibody (2). BCEs of protein antigens can be roughly classified into two categories, linear and conformational (3). Linear BCEs consist of residues that are contiguous in the antigen primary sequence, while the conformational BCEs comprise residues which are not contiguous in sequence but folding together in three-dimensional structure space. About 10% of BCEs are linear and about 90% are conformational (4). In our study, we focus on conformational BCEs of protein antigens.

The localization and identification of epitopes is of great importance for the development of vaccines and for the design of therapeutic antibodies (5, 6). However, traditional experimental methods to identify BCEs are still expensive and time-consuming (7). Therefore, great efforts for computational approaches based on machine learning algorithms have been developed to predict BCEs. These approaches can be divided in two categories: sequence-based and structure-based methods. As the name implies, the sequence-based approaches predict BCEs only based on the antigen sequence, while the structure-based approaches also consider its structural features. Currently, various structure-based predictors have been developed to predict and analyze BCEs including BeTop (8), Bpredictor (9), DiscoTope-2.0 (10), CE-KEG (11), CeePre (12), EpiPred (13), ASE Pred (14) and PECAN (15).

Some of those methods improve model performance by introducing novel features such as statistical features in BeTop, thick surface patch in Bpredictor, new spatial neighborhood definition and half-sphere exposure in DiscoTope-2.0, knowledge-based energy and geometrical neighboring residue contents in CE-KEG, B factor in CeePre and surface patches in ASE Pred. Except novel features, antibody structure information and suitable model also improve the performance of BCEs prediction. EpiPred utilizes antibody structure information to annotate the epitope region and improves global docking results. PECAN represents antigen or antibody structure as a graph and employ graph convolution operation on them to make aggregation of spatial neighboring residues. An additional attention layer is used to encode the context of the partner antibody in PECAN for predicting the more useful antibody-specific BCEs rather than antigenic residues. Because antibody structure information is required, these methods are not applicable to a novel virus when its antibody is unknown. However, all the currently structure-based BCEs prediction methods only use local information around target amino acid residue without considering the global information of the whole antigen sequence.

Global features have been proved to be effective in some biology sequence analysis models such as protein-protein interaction sites prediction model DeepPPISP (16) and protein phosphorylation sites prediction model DeepPSP (17). However, which model is used determines the effect of global features. DeepPPISP utilizes TextCNNs processing the whole protein sequence for protein-protein interaction sites prediction. DeepPSP employs SENet blocks and Bi-LSTM blocks to extract the global features for protein phosphorylation sites. In our study, we take advantage of the Attention based Bidirectional Long Short-Term Memory(Att-BLTM) networks. Att-BLTM networks are first introduced for relation classification in the field of natural language processing (NLP) (18). Att-BLSTM networks are also employed for some chemical and biomedical text processing tasks including chemical named entity recognition (19) and biomedical event extraction (20). Given the excellent performance of Att-BLSTM, we combine it with the novel deep learning model Graph Convolution Networks(GCNs) (21) for BCEs prediction.

In this study, we propose a structure-based BCEs prediction model utilizing both antigen local features and global features. By combining Att-BLSTM and GCNs, both local and global features are used in our model to improve its prediction performance. We implement our model on some public datasets and the results show that global features can provide useful information for BCEs prediction.

## 2 MATERIALS AND METHODS

### 2.1 Datasets

In order to make fair comparison, we use the same antibody-antigen complexes as PECAN (15). It should be noted that those bound conformations are only uesd to identify epitope residues and no-epitope residues. Same as previous works (13, 15), residues are labeled as part of the BCEs if they have any heavy atom within 4.5Å away from any heavy atom on the antibody. As our model is partner independent, it only takes antigen structure as input for predicting BCEs.

Those complexes are from two separate datasets: EpiPred (13) and Docking Benchmarking Dataset (DBD) v5 (22). The 148 antibody-antigen complexes from EpiPred share no more than 90% pairwise sequence identity. Among them, 118 complexes are used for training and 30 for testing. For constructing a separate validation set, PECAN filters the antibody-antigen complexes in DBD v5 and selects 162 complexes which have no more than 25% pairwise sequence identity to every antigen in the testing set. Antigens in training set are used for training our model, antigens in valdation set are used to tune the hyperparameters of our proposed method, and antigens in testing set are uesd for evaluation our model and making comparision with competing methods. The size of datasets and number of BCEs are shown in Table 1.

**Table 1.**
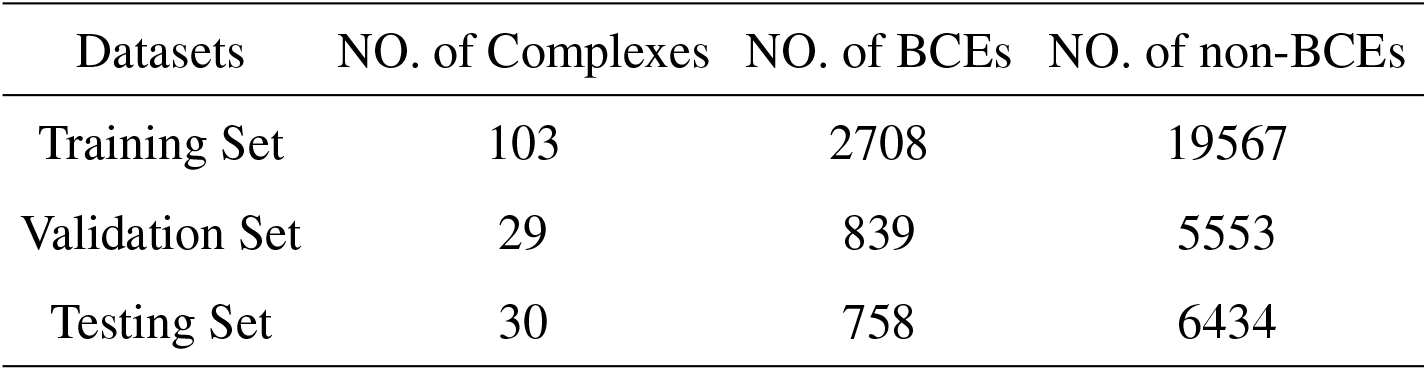
Summary of datasets.

### 2.2 Input features representation

For global features, we construct the input antigen sequence as a set of sequential residues:

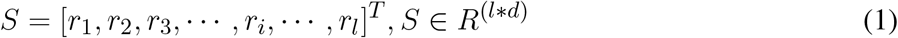

where each residue is represented as a vector *r*_*i*_ ∈*R*^*d*^ corresponding to the *i*-th residue in the antigen sequence, *l* is the antigen sequence length, and *d* is the residue feature dimension.

For local features, each antigen structure is represented as a graph as related studies (13, 15). The residue is a node in the protein graph whose features represent its properties. For residue *r*_*i*_, the local environment *N*_*i*_ consists of *k* spatial neighboring residues:

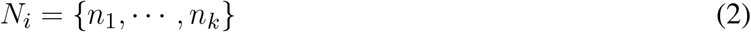

And, 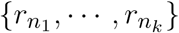 are the neighbors of residue *r*_*i*_ which define the operation field of the graph convolution.

In this study, node features and edge features in antigen graph are used for characterizing the local environment of target residue. The node features are represented as a 128-dimension vector encoding important properties as in our earlier work (23). All those node features can be divided into two classes: sequence-based and structure-based. Sequence-based features consist of the one-hot encoding of the amino acid residue type, seven physicochemical parameters (24) and a conservation profile returned by running PSI-BLAST (25) against nr database (26). The structure-based features are calculated for each antigen structure isolated from the antibody-antigen complex by DSSP (27), MSMS (28), PSAIA (29) and Biopython (30). The edge features between two residues *r*_*i*_ and *r*_*j*_ are representing as *e*_*ij*_. *e*_*ij*_ reflects the spatial relationships including the distance and angle between residue pair *r*_*i*_ and *r*_*j*_ and it is computed by their *C*_*a*_ (31).

### 2.3 Model architecture

Our model solves a binary classification problem: judging an antigen residue binding to antibody or not. As shown in Figure 1, our model consists of two parallel parts: GCNs and Att-BLSTM networks. The former captures local features of target antigen residue from its spatial neighbors by using graph convolutional layer, and the latter extracts global features from the whole antigen sequence by using Bi-LSTM layer and attention layer. The outputs are concatenated and fed to fully connected layer to predict the binding probability for each antigen residue.

**Figure 1.**
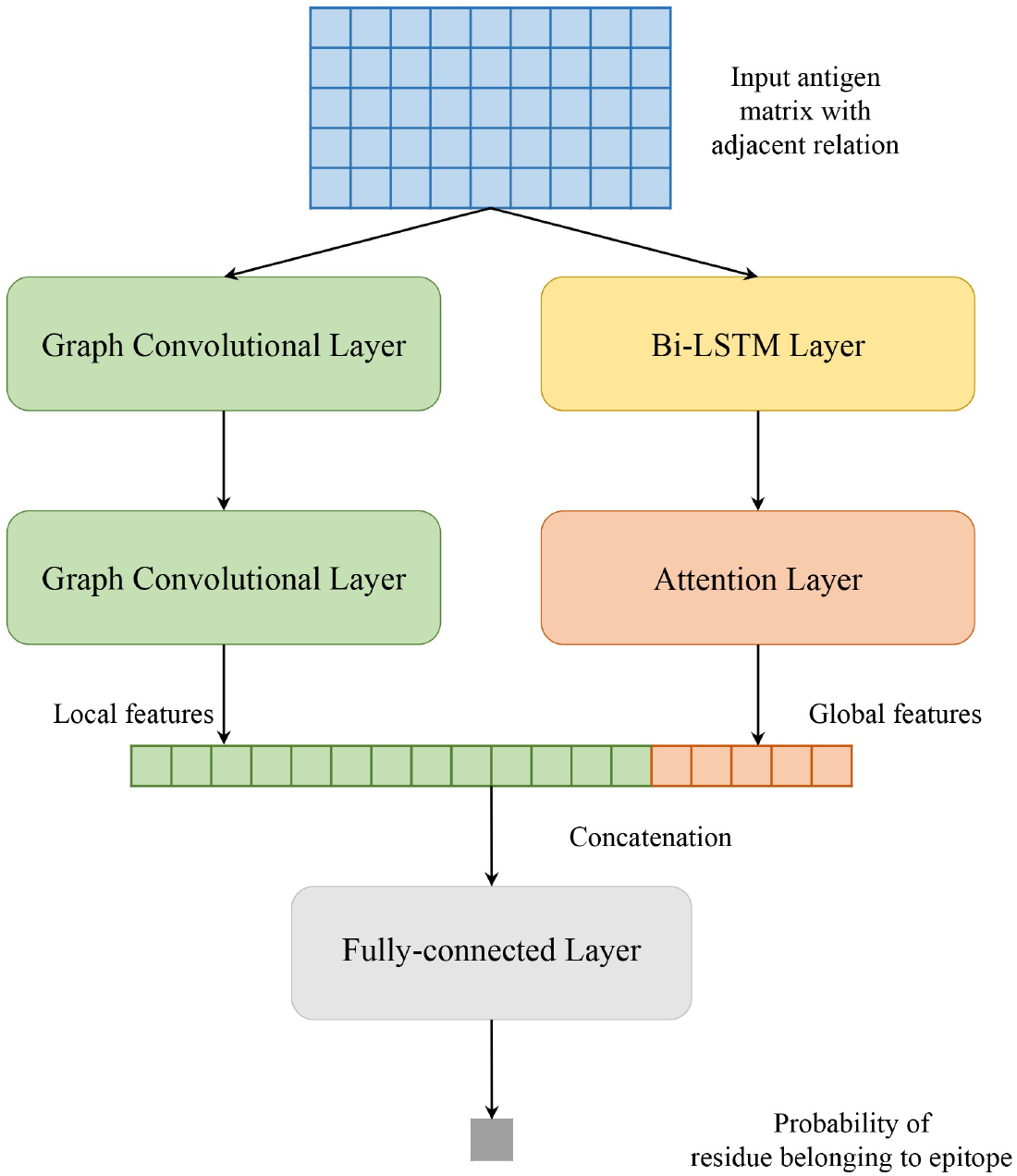
Model architecture of proposed method in this study.

#### 2.3.1 Graph convolutional networks

Figure 2 shows the flow of convolution operation using the information of nodes and edges. At first, each protein is represented as a graph, and a residue is a node in the graph. The local environment of the target residue is a set of residues which are adjacent in space. And then, node and edge are represented by a vector as our previous work (23). Actually, the graph convolution operation on the local environment of target residue is the aggregation of neighboring residues and its edges. Every node in the graph is updated through repeated aggregation operation. Based on edges are used or not, we utilize two graph convolution operators in this study:

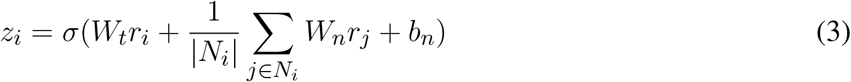

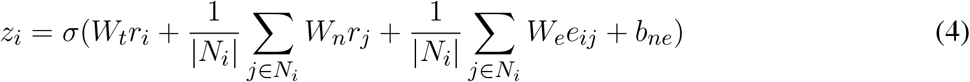

where *N*_*i*_ is the receptive field, i.e. a set of neighbors of target residue *r*_*i*_, *W*_*t*_ is the weight matrix associated with the target node, *W*_*n*_ is the weight matrix associated with neighboring nodes, *σ* is a non-linear activation function, and *b*_*n*_ is a bias vector. Formula 3 groups the node information in receptive filed. Formula 4 utilizes not only node features but also edge feature between two residues, where *W*_*e*_ is the weight matrix associated with edge feature, *e*_*ij*_ represents the edge feature between residue *r*_*i*_ and *r*_*j*_, and *b*_*ne*_ is a vector of biases.

**Figure 2.**
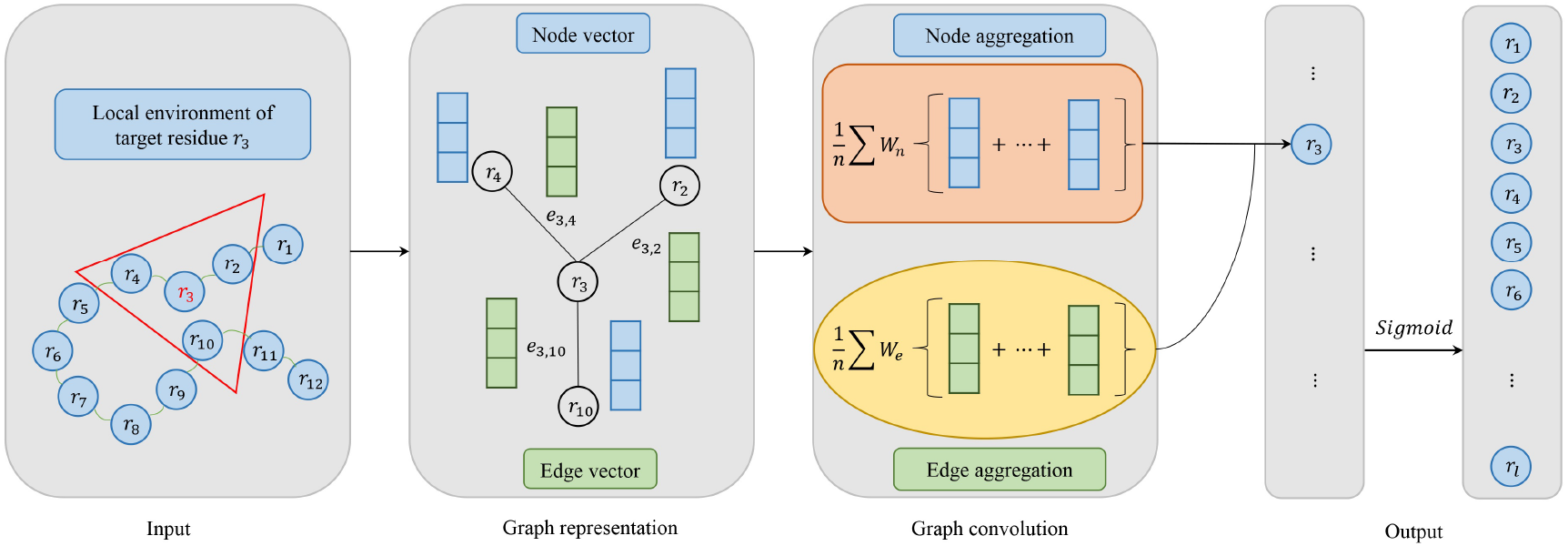
Model architecture of graph convolutional networks consisting of four parts: input, graph representation, graph convolution and output. In the input, the target residue is red and the local environment of the target residue is in a red triangle. For graph representation, node vector is blue and edge vector is green. For graph convolution, if edge vector is used, it corresponds to the formula 4. Otherwise, it corresponds to the formula 3. In the output, we use the Sigmoid activation function.

#### 2.3.2 Attention-based Bidirectional Long Short-Term Memory networks

Besides local features, global features are crucial in BCEs prediction as well. In our work, Attention-based Bidirectional Long Short-Term Memory(Att-BLSTM) networks are used to capture global sequence information of input antigen sequence. Currently, Att-BLSTM has been used for processing chemical and biomedical text (19, 20). It can capture the most important semantic information in a sequence. However, its advantage has not been exploited in biology sequence analysis such as BCEs prediction.

Figure 3 shows the architecture of Att-BLSTM. At first, the input antigen matrix *S* is fed into a Long Short-Term Memory (LSTM) network which learns long-range dependencies in a sequence (32). Typically, the structure of an LSTM unit at each time t is calculated by the following formulas:

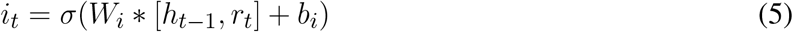

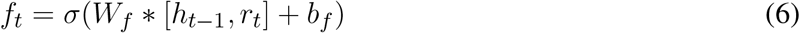

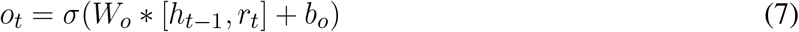

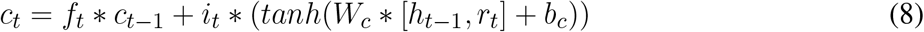

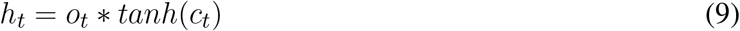

where *tanh* is the element-wise hyperbolic tangent, *σ* is the logistic sigmoid function, *r*_*t*_, *h*_*t−*1_ and *c*_*t−*1_ are inputs, and *h*_*t*_ and *c*_*t*_ are outputs. There are three gates consisting of one input gate *i*_*t*_ with corresponding weight matrix *W*_*i*_, and a bias *b*_*i*_; one forget gate *f*_*t*_ with corresponding weight matrix *W*_*f*_, and a bias *b*_*f*_ ; one output gate *o*_*t*_ with corresponding weight matrix *W*_*o*_, and a bias *b*_*o*_.

**Figure 3.**
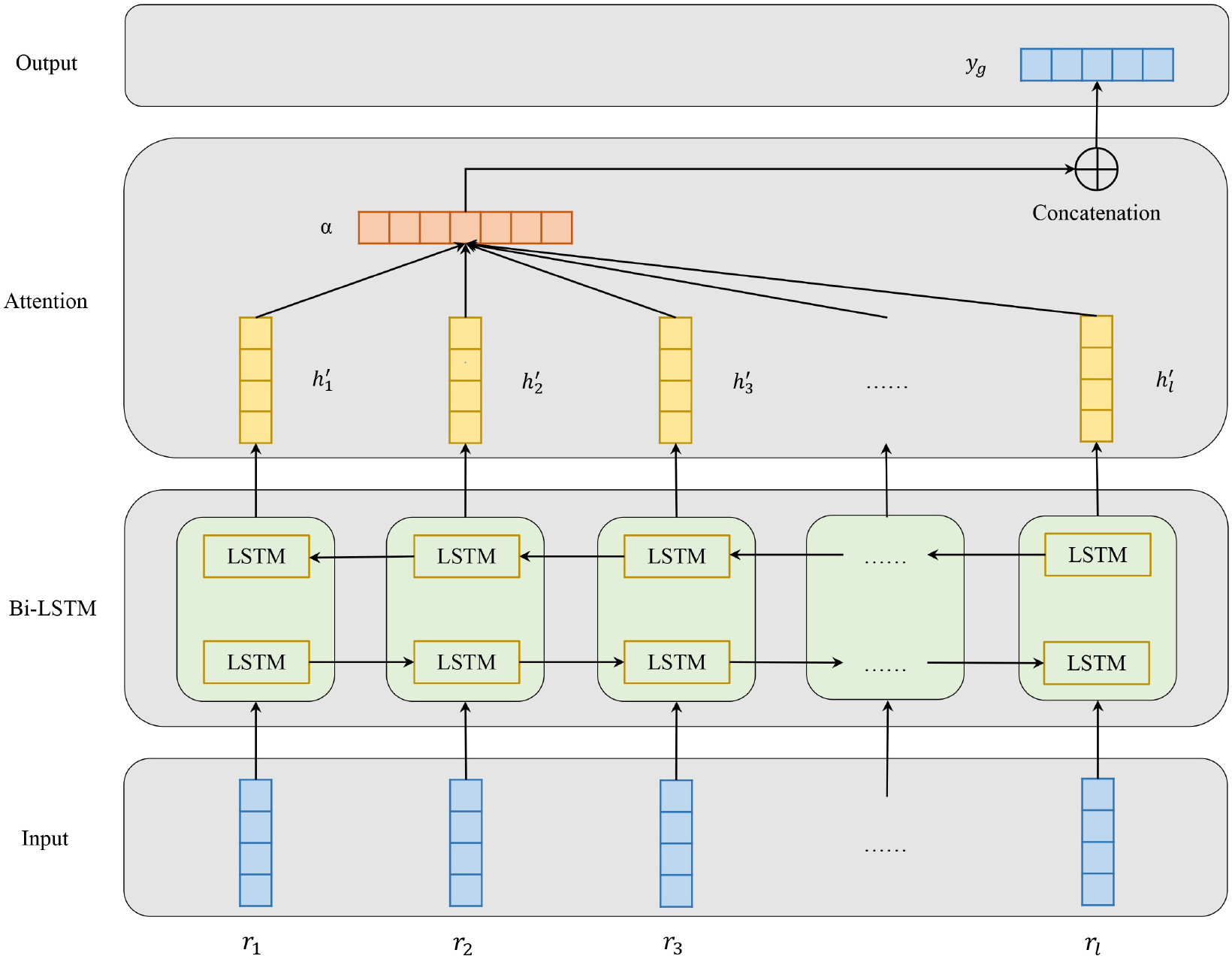
Model architecture of attention-based bidirectional long short-term memory networks which consists of four parts: input, Bi-LSTM layer, Attention layer and output.

Bidirectional LSTM(Bi-LSTM) can learn forward and backward information of input sequence. As shown in Figure 2, the networks contain two sub-networks for the left and right sequence contexts. For the *i*-th residue in the input antigen sequence, we combine the forward pass output 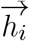 and backward pass output 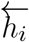 by concatenating them:

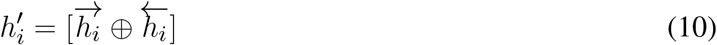

The output of Bi-LSTM layer is matrix H which consists of all output vectors of input antigen residues: 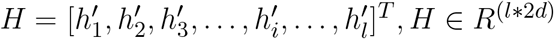, where *l* is the input antigen sequence length, and *d* is the residue features dimension.

Attention mechanism has been used in a lot of biology tasks ranging from compound-protein interaction prediction (33), paratope prediction (15) and protein structure prediction (34). The attention layer in our model employs a classical additive model in which *α* is the attention weight. After attention layer of Att-BLSTM, the novel representation *S′* as well as the output *y*_*g*_ of the input antigen is formed by a weighted sum of those output vectors *H*:

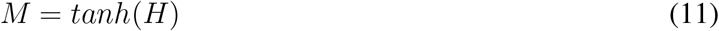

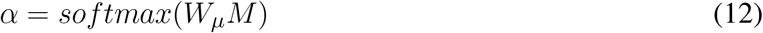

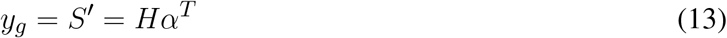

#### 2.3.3 Fully-connected networks

As shown in Figure 1, the local features *z*_*i*_ extracted by GCNs and the global features *y*_*g*_ derived from Att-BLSTM networks are concatenated. And then, they are fed to fully-connected layer. The calculation of probability *y*_*i*_ for each input antigen residue belonging to epitope is shown as:

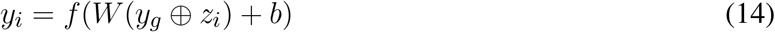

### 2.4 Performance evaluation

In order to make comparison with state-of-the-art structure-based BCEs predictors, we use three evaluation metrics to evaluate the performances of the BCEs prediction models: Precision, Recall and Matthews Correlation Coefficient (MCC) which are shown as followings:

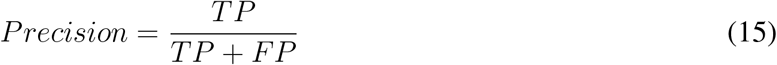

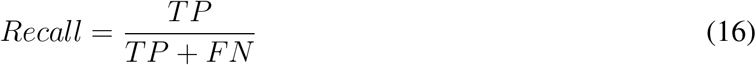

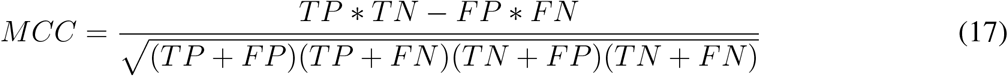

where, TP(True Positive) is the number of interacting residues that are correctly predicted as BCEs, FP(False Positive) is the number of non-interacting residues that are falsely predicted as BCEs, TN(True Negative) denotes the number of non-interacting sites that are identified correctly, and FN(False Negative) denotes the number of interacting sites that are identified falsely. Precision and recall reflect the prediction tendencies of classifiers. Recall indicates the percentage of correct predictions for positive and negative samples. Precision shows the percentage of correct positive samples. There is a trade-off between precision and recall. Recall favors positive-bias predictions, while precision favors negative predictions.

Because precision, recall and MCC are threshold-dependent, we also utilize the area under the receiver operating characteristics curve (AUC ROC) and precision-recall curve(AUC PR) which gives a threshold-independent evaluation on the overall performance. Moreover, AUC PR is more sensitive than AUC ROC on imbalanced data (35). And, the datasets used for BCEs prediction are roughly 90% negative class. Therefore, we take AUC PR as the most import metric for model evaluation and selection.

It should be noted that the precision, recall and MCC shown in Table 2 are averaged over all antigens in the testing set. And, the AUC ROC and AUC PR reported in Figure 4 and 5 are calculated among all antigen residues in the testing set.

**Table 2.**
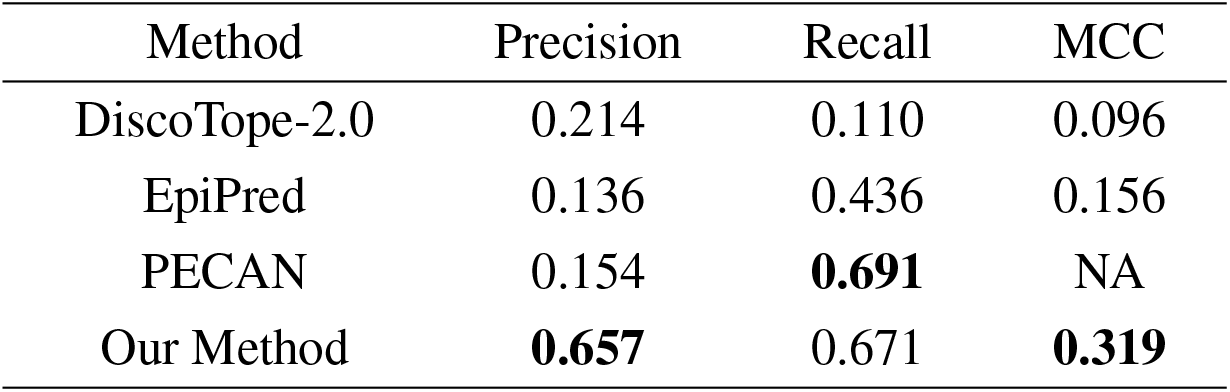
Performances of BCEs prediction methods.

**Figure 4.**
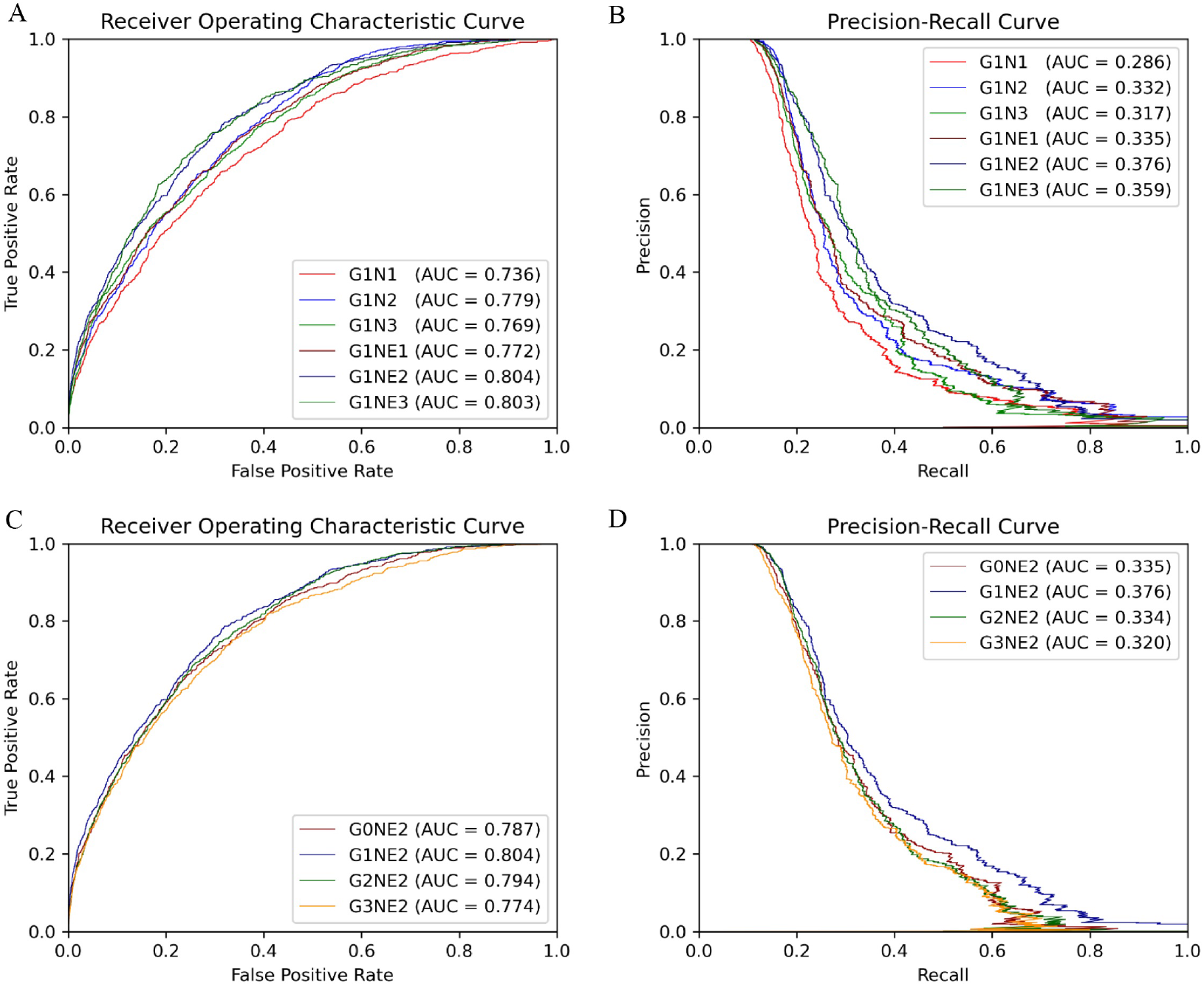
ROC and PR curve of different network combinations among all antigen residues in the testing set.

**Figure 5.**
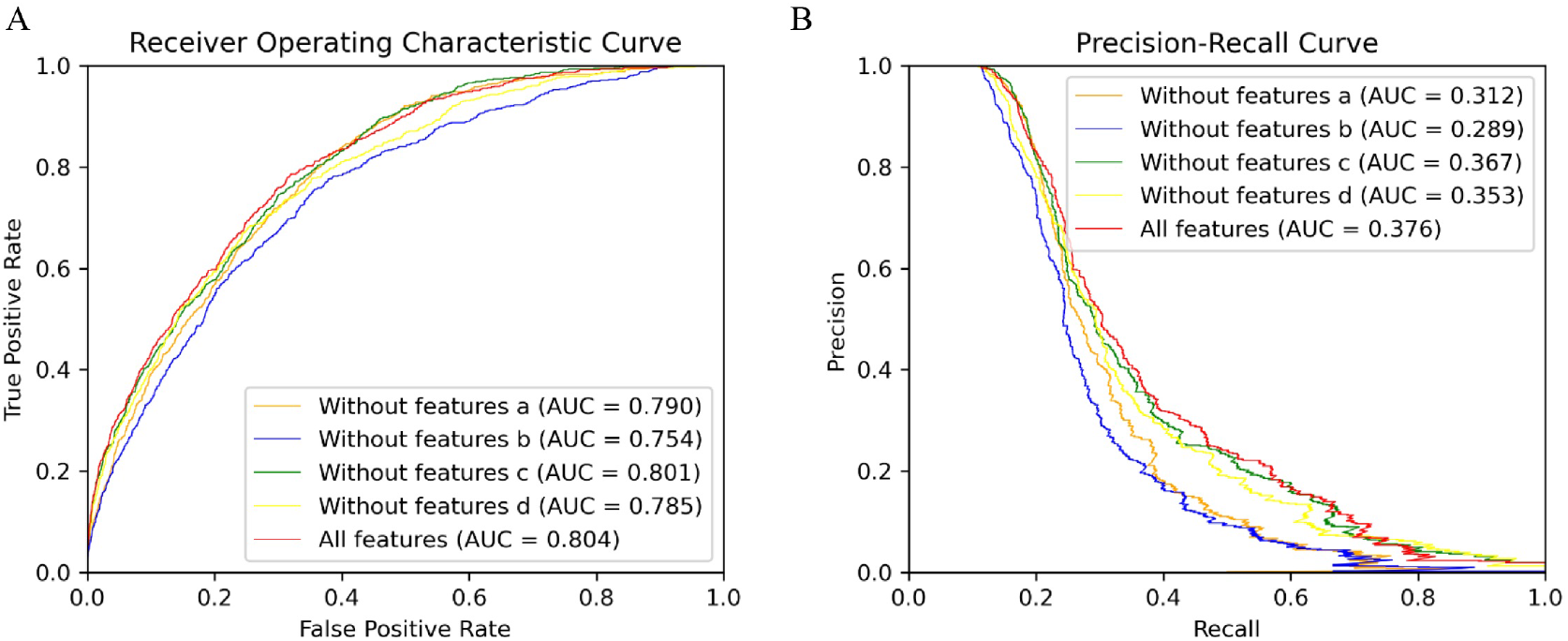
ROC and PR curve of different combinations of input features among all antigen residues in the testing set.

### 2.5 Implementation details

We implement our model using PyTorch. The training details of these neural networks are as follows: optimization: Momentum optimizer with Nesterov accelerated gradients; learning rate: 0.1, 0.01, 0.001 and 0.0001; batch size: 32, 64 and 128; dropout: 0.2, 0.5 and 0.7; spatial neighbors in the graph: 20; number of LSTM layers in Att-BLSTM networks: 1, 2 or 3; number of graph convolution networks layers: 1, 2 or 3. Training time of each epoch varies from roughly 1 to 3 minutes depending on network depth, using a single NVIDIA RTX2080 GPU.

For each combination, networks are trained until the performance on the validation set stops improving or for a maximum of 250 epochs. Graph convolution networks have the following number of filters for 1, 2 and 3 layers, respectively: (256), (256, 512), (256, 256, 512). All weight matrices are initialized as (31) and biases are set to zero.

## 3 RESULTS AND DISCUSSION

### 3.1 The effects of different network combinations

In this section, we focus on which network combinations are most effective. The AUC ROC and AUC PR are shown in Figure 4, where label G denotes the model processing global features and the number after G indicates the model depth, for instance, G1 represents 1-layer Att-BLSTM. And N denotes residue Node features and NE denotes both residue Node and Edge features are used in GCNs. The number behind N or NE indicates the depth of GCNs. For example, G1NE2 represents model with 1-layer Att-BLSTM and 2-layer GCNs using residue node and edge features.

First, we train our model of 1-layer Att-BLSTM with varying GCNs depths with or without residue edge features. From Figure 4 A and B, we observe that the 2-layer GCNs with residue edge features perform best (AUC ROC = 0.804, AUC PR = 0.376). This draws the same conclusion with our earlier work for antibody paratope prediction (23). We also find that residue edge features can always provide better performance as the GCNs depths vary. The same results are found in protein interface prediction task using GCNs as well (31).

Second, 2-layer GCNs and Att-BLSTM networks of different depths are combined in our model. Figure 4 C and D show the performance evaluated by AUC ROC and AUC PR. It can be found that the combination of 1-layer Att-BLSTM network and 2-layer GCNs with residue edge features still has the best results. In general, the deeper the Att-BLSTM networks grow, the results get worse. As discussed in DeepPPISP (16), global features may cover the relationships among residues of longer distances. However, those relationships might become weaken when the Att-BLSTM networks grow deeper.

In summary, our model with 2-layes GCNs and 1-layer Att-BLSTM network performs best, and it is the proposed model in this paper and used for comparison with competing methods in the following sections.

### 3.2 The effects of global features

The global feature has been shown to improve the performance of protein-protein interaction sites prediction in DeepPPISP (16) and protein phosphorylation sites prediction in DeepPSP (17). In order to verify whether global features are effective in BCEs prediction as well, we remove the Att-BLSTM networks in our model for comparison. As shown in Figure 4 C and D, label G0 means there is only GCNs in our model, and no global features are used. Without global features, the AUC ROC is 0.787, which is lower than the proposed model G1NE2 (also lower thanG2NE2, but slightly better than G3NE2). Without global features, the AUC PR is 0.335, which is significantly worse than the proposed model (but slightly better than G2NE2 and G3NE2). The model without global features performs worse on both AUC ROC and AUC PR metrics than our proposed model. Therefore, global features improve the performance of our model for BCEs prediction.

However, models with global features not always outperform model without global features in our experiment. Similar observation has been found in DeepPSP, but DeepPPISP reaches a contrary conclusion. This situation might be caused by different models processing global features. In DeepPPISP, a simple fully-connected network is used, and in DeepPSP, SENet blocks and Bi-LSTM blocks are used.

### 3.3 The effects of different types of input features

Different types of input features (sequence and structure-based) play different roles in our model. The input features consist of four types: residue type one-hot encoding (a), profile features (b), seven physicochemical parameters (c) and structural features (d). To discover what role each feature type plays in our method, we delete each input feature type and compare their performances on our proposed model (G1NE2, i.e. 1-layer Att-BLSTM network and 2-layersGCNs with residue edge features). Figure 4 shows the experimental results. As Figure 5 shows, the AUC ROC without features b is 0.754, significantly lower than the best performance 0.804. The AUC PR without features b drops biggest from 0.376 (all features) to 0.289. It indicates that profile features (feature type b) are most important in our model for BCEs prediction. The model using all the features still performs best on both AUC ROC and AUC PR metrics.

### 3.4 Comparison with competing methods

To evaluate the performance of our method for BCEs prediction, we compare our proposed model with three competing structure-based BCEs prediction methods: DiscoTope-2.0 (10), EpiPred (13) and PECAN (15). Note that these methods all used local features but did not consider global features. The precision, recall and MCC calculated in this study using a threshold 0.116 at which our method achieve best performance on the testing set. Table 2 shows the experimental results of our method and the competing models. The results on three competing models are taken from (13). Although our model gets lower Recall than PECAN, it is higher than all other competing methods on Precision and MCC.

We also compare the results of each antigen in testing set with DiscoTope-2.0 and EpiPred. The results presented of DiscoTope-2.0 and EpiPred in Table S1 are taken form (13). The values in bold indicate the best prediction result. We find that our model achieves best Precision on 26 antigens, best Recall on 20 antigens and best MCC on 20 antigens of all 30 antigens in testing set. We also observe that our model produces usable prediction even for the long antigen target as the global features provide information from long distance effect.

### 3.5 Case study

Our method is also applied to the structure data (PDB 7BWJ) of SARS-Cov-2 Receptor Binding Domain (RBD) in the complex with an antibody Fab (P2B-2F6). The entry of SARS-CoV-2 into its target cells depends on binding between the RBD of the viral spike protein and its cellular receptor, angiotensin-converting enzyme 2 (ACE2) (36). P2B-2F6 can not only prevent the RBD from binding to ACE2 through steric hindrance but also can compete with ACE2 to bind to the RBD. As Figure 6 shows, the binding interfaces are overlapping at G466 and Y499 (37) which are in green.

**Figure 6.**
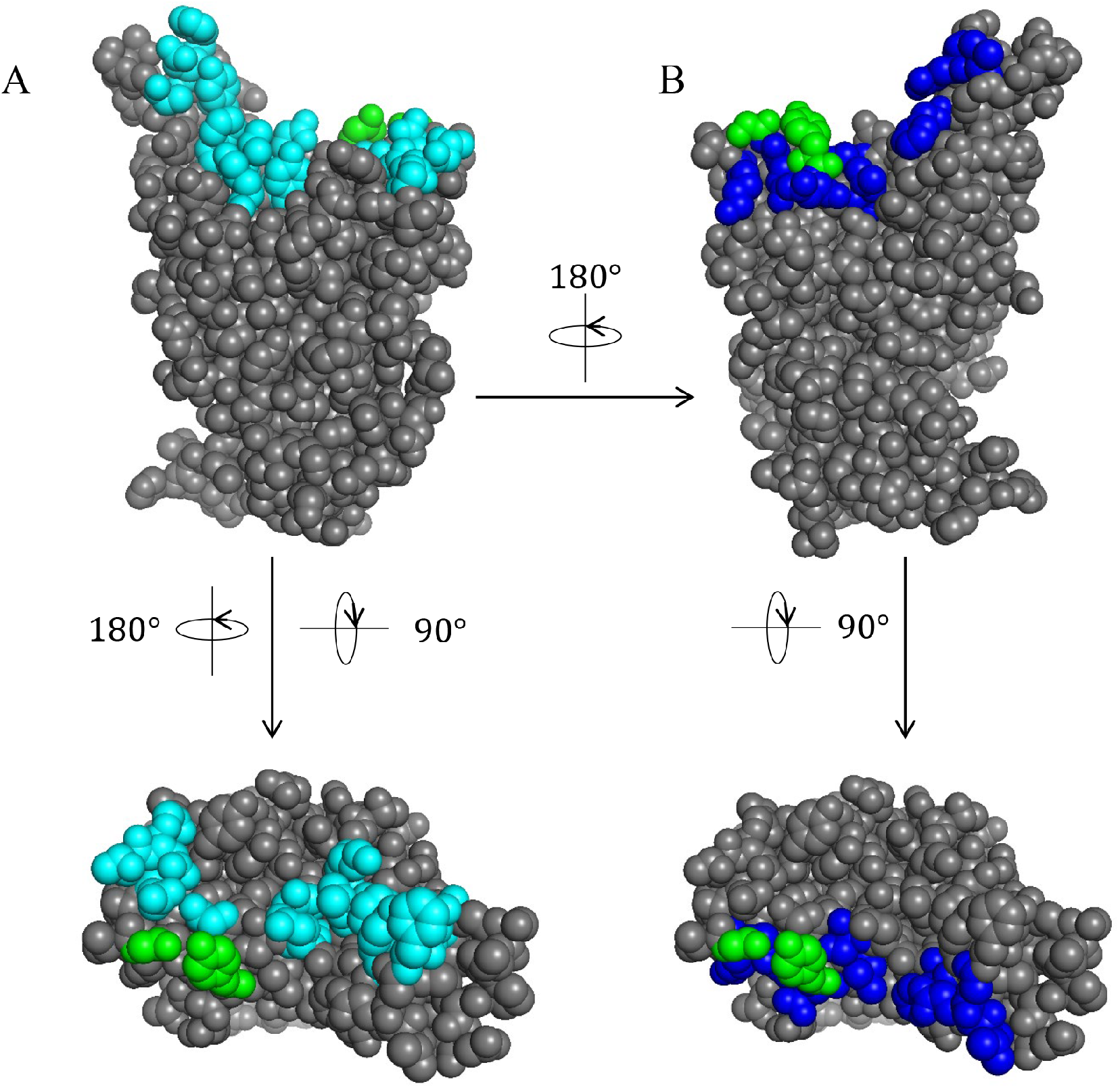
SARS-Cov-2 RBD binding interface with ACE2 and P2B-2F6. (A) The binding interface with ACE2 is cyan. (B) The binding interface with P2B-2F6 is blue. (A-B) The overlapping residues recognized by ACE2 and P2B-2F6 are in green.

The BCEs prediction results are listed in Table 3. Compared with the competing predictors, our method achieves the best performance for every metric. The precision of our method is highest, which indicates that more predicted BCEs are true BCEs. There is a significant improvement in recall, which implies that more true BCEs are identified by our method. For MCC, which metric characterizes the overall performance of classifiers, our method outperforms all other competing predictors.

**Table 3.** Prediction performances on the SARS-Cov-2 RBD.

| Method | Precision | Recall | MCC |
| --- | --- | --- | --- |
| DiscoTope-2.0 | 0.598 | 0.695 | 0.276 |
| EpiPred | 0.483 | 0.462 | -0.051 |
| PECAN | 0.523 | 0.551 | 0.068 |
| Our Method | **0.636** | **0.815** | **0.414** |

It can be seen from Figure 7 and Table 3, our method identifiesthe most true BCEs among all predictors. However, some non-BCEs are also corrected as true BCEs by our method. Compared with the state-of-the-art method PECAN (15), our method has an impressive clustering effect on true BCEs.

**Figure 7.**
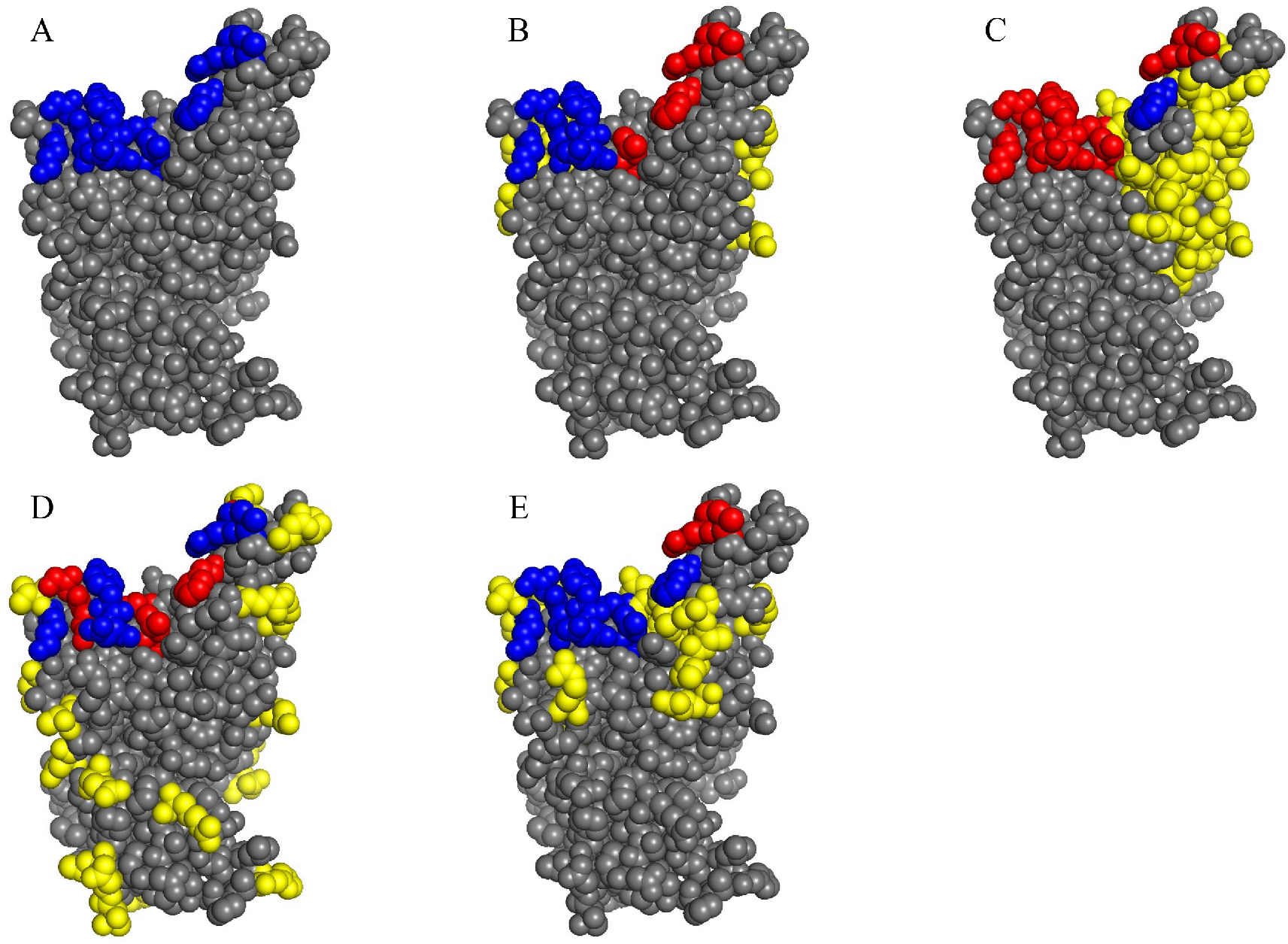
BCEs prediction for SARS-Cov-2 RBD (PDB 7BWJ). (A) The true epitope residues. (B-E) Prediction results by Discotope-2.0, EpiPred, PECAN and our method, respectively. TP predictions are in blue, FN predictions are in red, FP predictions are in yellow and the background grey represents TN predictions.

## 4 CONCLUSIONS

Accurate prediction of BCEs is helpful for understanding the basis of immune interaction and is beneficial to therapeutic design. In this work, we propose a novel deep learning framework combining local and global features which are extracted from antigen sequence and structure to predict BCEs. GCNs are used for capturing the local features of a target residue. Att-BLSTM networks are used to extract global features, which figure the relationship between a target residue and the whole antigen. We employ our model on a public and popular dataset and the results show improvement of BCEs prediction. Moreover, our results declare that the global features are useful for improving the prediction of BCEs.

Though our method outperforms other competing computational methods for BCEs prediction, it also has some disadvantages. The first one is that our predictor needs antigen structure as it takes structure-based residue features as input. The second one is that it consumes long computer time because PSI-BLAST (25) needs to be performed at the stage of extracting residue features.

For deep case study, we apply our method to the BCEs prediction for SARS-Cov-2 RBD. It has been found that our method not only provides more accurate predictions of BCEs but also shows more meaningful BCEs prediction by clustering adjacent residues and utilizing long distance effect.

In this study, we show that combing local and global features can be useful for BCEs prediction. In the future, we would further improve BCEs prediction by expanding the training set.

## AUTHOR CONTRIBUTIONS

SL designed the study. SL and YL performed the method development. SL, XN and QM performed the data analysis. XN and SZ wrote and revised the manscript. All authors reviewed the manuscript.

## REFERENCES

1. Getzoff ED, Tainer JA, Lerner RA, Geysen HM. The Chemistry and Mechanism of Antibody Binding to Protein Antigens. Advances in Immunology 43 (1988) 1–98. doi:10.1016/S0065-2776(08)60363-6.

2. Michnick SW, Sidhu SS. Submitting antibodies to binding arbitration. Nature Chemical Biology 4 (2008) 326–329. doi:10.1038/nchembio0608-326.

3. Barlow DJ, Edwards MS, Thornton JM. Continuous and discontinuous protein antigenic determinants. Nature 322 (1986) 747–748. doi:10.1038/322747a0.

4. Caoili SEC. Hybrid methods for B-cell epitope prediction approaches to the development and utilization of computational tools for practical applications. Methods in Molecular Biology 1184 (2014) 245–283. doi:10.1007/978-1-4939-1115-814.

5. Gershoni JM, Roitburd-Berman A, Siman-Tov DD, Freund NT, Weiss Y. Epitope mapping: The first step in developing epitope-based vaccines. BioDrugs 21 (2007) 145–156. doi:10.2165/00063030-200721030-00002.

6. Chan AC, Carter PJ. Therapeutic antibodies for autoimmunity and inflammation. Nature Reviews Immunology 10 (2010) 301–316. doi:10.1038/nri2761.

7. Abbott WM, Damschroder MM, Lowe DC. Current approaches to fine mapping of antigen-antibody interactions. Immunology 142 (2014) 526–535. doi:10.1111/imm.12284.

8. Zhao L, Wong L, Lu L, Hoi SC, Li J. B-cell epitope prediction through a graph model. BMC Bioinformatics 13 (2012) 1–12. doi:10.1186/1471-2105-13-S17-S20.

9. Zhang W, Xiong Y, Zhao M, Zou H, Ye X, Liu J. Prediction of conformational B-cell epitopes from 3D structures by random forests with a distance-based feature. BMC Bioinformatics 12 (2011) 1–10. doi:10.1186/1471-2105-12-341.

10. Kringelum JV, Lundegaard C, Lund O, Nielsen M. Reliable B Cell Epitope Predictions: Impacts of Method Development and Improved Benchmarking. PLoS Computational Biology 8 (2012) e1002829. doi:10.1371/journal.pcbi.1002829.

11. Lo YT, Pai TW, Wu WK, Chang HT. Prediction of conformational epitopes with the use of a knowledge-based energy function and geometrically related neighboring residue characteristics. BMC Bioinformatics 14 (2013) 1–10. doi:10.1186/1471-2105-14-S4-S3.

12. Ren J, Liu Q, Ellis J, Li J. Tertiary structure-based prediction of conformational B-cell epitopes through B factors. Bioinformatics 30 (2014) 264–273. doi:10.1093/bioinformatics/btu281.

13. Krawczyk K, Liu X, Baker T, Shi J, Deane CM. Improving B-cell epitope prediction and its application to global antibody-antigen docking. Bioinformatics 30 (2014) 2288–2294. doi:10.1093/bioinformatics/btu190.

14. Jespersen MC, Mahajan S, Peters B, Nielsen M. Antibody Specific B-Cell Epitope Predictions : Leveraging Information From Antibody-Antigen Protein Complexes. Frontiers in Immunology 10 (2019) 1–10. doi:10.3389/fimmu.2019.00298.

15. Pittala S, Bailey-Kellogg C. Learning context-aware structural representations to predict antigen and antibody binding interfaces. Bioinformatics 36 (2020) 3996–4003. doi:10.1093/bioinformatics/btaa263.

16. Zeng M, Zhang F, Wu FX, Li Y, Wang J, Li M. Protein-protein interaction site prediction through combining local and global features with deep neural networks. Bioinformatics 36 (2020) 1114–1120. doi:10.1093/bioinformatics/btz699.

17. Guo L, Wang Y, Xu X, Cheng KK, Long Y, Xu J, et al. DeepPSP: A Global-Local Information-Based Deep Neural Network for the Prediction of Protein Phosphorylation Sites. Journal of Proteome Research 20 (2021) 346–356. doi:10.1021/acs.jproteome.0c00431.

18. Zhou P, Shi W, Tian J, Qi Z, Li B, Hao H, et al. Attention-based bidirectional long short-term memory networks for relation classification. Annual Meeting of the Association for Computational Linguistics (2016), 207–212. doi:10.18653/v1/p16-2034.

19. Luo L, Yang Z, Yang P, Zhang Y, Wang L, Lin H, et al. An attention-based BiLSTM-CRF approach to document-level chemical named entity recognition. Bioinformatics 34 (2018) 1381–1388. doi:10.1093/bioinformatics/btx761.

20. Luo L, Yang Z, Yang P, Zhang Y, Wang L, Lin H, et al. An attention-based BiLSTM-CRF approach to document-level chemical named entity recognition. Bioinformatics 34 (2018) 1381–1388. doi:10.1093/bioinformatics/btx761.

21. Kipf. Semi-supervised classfication with graph convolutional networks. International Conference on Learning Representations (2017), 1–10.

22. Vreven T, Moal IH, Vangone A, Pierce BG, Kastritis PL, Torchala M, et al. Updates to the Integrated Protein-Protein Interaction Benchmarks: Docking Benchmark Version 5 and Affinity Benchmark Version 2. Journal of Molecular Biology 427 (2015) 3031–3041. doi:10.1016/j.jmb.2015.07.016.

23. Lu S, Li Y, Wang F, Nan X, Zhang S. Leveraging Sequential and Spatial Neighbors Information by Using CNNs Linked With GCNs for Paratope Prediction. IEEE/ACM Transactions on Computational Biology and Bioinformatics 19 (2022) 68–74. doi:10.1109/TCBB.2021.3083001.

24. Meiler J, Müller M, Zeidler A, Schmäschke F. Generation and evaluation of dimension-reduced amino acid parameter representations by artificial neural networks. Journal of Molecular Modeling 7 (2001) 360–369. doi:10.1007/s008940100038.

25. Altschul SF, Madden TL, Schäffer AA, Zhang J, Zhang Z, Miller W, et al. Gapped BLAST and PSI-BLAST: a new generation of protein database search programs. Nucleic Acids Research 25 (1997) 3389–3402.

26. McGinnis S, Madden TL. BLAST: At the core of a powerful and diverse set of sequence analysis tools. Nucleic Acids Research 32 (2004) 20–25. doi:10.1093/nar/gkh435.

27. Kabsch W, Sander C. Dictionary of protein secondary structure: Pattern recognition of hydrogen-bonded and geometrical features. Biopolymers 22 (1983) 2577–2637. doi:10.1002/bip.360221211.

28. Sanner MF, Olson AJ, Spehner J. Reduced surface: An efficient way to compute molecular surfaces. Biopolymers 38 (1996) 305–320. doi:10.1002/(sici)1097-0282(199603)38:3⟨305::aid-bip4⟩3.3.co;2-8.

29. Mihel J, Šikić M, Tomić S, Jeren B, Vlahoviček K. PSAIA - Protein structure and interaction analyzer. BMC Structural Biology 8 (2008) 1–11. doi:10.1186/1472-6807-8-21.

30. Cock PJ, Antao T, Chang JT, Chapman BA, Cox CJ, Dalke A, et al. Biopython: Freely available Python tools for computational molecular biology and bioinformatics. Bioinformatics 25 (2009) 1422–1423. doi:10.1093/bioinformatics/btp163.

31. Fout A, Byrd J, Shariat B, Ben-Hur A. Protein interface prediction using graph convolutional networks. Conference on Neural Information Processing Systems (2017), 6531–6540.

32. Hochreiter S, Schmidhuber J. Long Short-Term Memory. Neural Computation 9 (1997) 1735–1780.

33. Chen L, Tan X, Wang D, Zhong F, Liu X, Yang T, et al. TransformerCPI: Improving compound–protein interaction prediction by sequence-based deep learning with self-attention mechanism and label reversal experiments. Bioinformatics 36 (2020) 4406–4414. doi:10.1093/bioinformatics/btaa524.

34. Berkeley UC, Meier J, Sercu T, Rives A. Transformer Protein Language Models Are Unsupervised Structure Learners. bioRxiv (2020) 1–24.

35. Staeheli LA, Mitchell D. The Relationship Between Precision-Recall and ROC Curves Jesse. International Conference on Machine Learning (2006), 233–240. doi:10.1145/1143844.1143874.

36. Zhou P, Yang XL, Wang XG, Hu B, Zhang L, Zhang W, et al. A pneumonia outbreak associated with a new coronavirus of probable bat origin. Nature 579 (2020) 270–273. doi:10.1038/s41586-020-2012-7.

37. Lan J, Ge J, Yu J, Shan S, Zhou H, Fan S, et al. Structure of the SARS-CoV-2 spike receptor-binding domain bound to the ACE2 receptor. Nature 581 (2020) 215–220. doi:10.1038/s41586-020-2180-5.

