## Supplementary material for "A Structure-based B-cell Epitope Prediction Model Through Combing Local and Global Features": Table S1

| PDB ID | Ag Size | DiscoTope-2.0 | | | EpiPred | | | Our model | | |
|---|---|---|---|---|---|---|---|---|---|---|
| | | Precision | Recall | MCC | Precision | Recall | MCC | Precision | Recall | MCC |
| 4hj0 | 92 | 0 | 0 | 0 | 0.32 | **0.9** | **0.27** | **0.61** | 0.621 | 0.231 |
| 1tzh | 94 | 0.73 | **0.87** | **0.72** | 0.01 | 0.06 | 0.04 | **0.692** | 0.683 | 0.376 |
| 4am0 | 96 | 0.33 | 0.2 | **0.19** | 0.13 | **0.7** | 0.09 | **0.547** | 0.569 | 0.114 |
| 2ih3 | 97 | 0 | 0 | 0 | 0.16 | 0.64 | 0.08 | **0.755** | **0.765** | **0.519** |
| 4i77 | 97 | 0 | 0 | 0 | 0.23 | 0.55 | 0 | **0.807** | **0.777** | **0.584** |
| 3q1s | 113 | 0 | 0 | 0 | 0.19 | 0.81 | 0.15 | **0.673** | **0.754** | **0.419** |
| 1p2c | 129 | **1** | 0.05 | 0 | 0 | 0 | 0 | 0.68 | **0.806** | **0.469** |
| 4ht1 | 131 | 0 | 0 | 0 | 0.05 | 0.14 | 0.05 | **0.622** | **0.59** | **0.21** |
| 3ab0 | 136 | 0 | 0 | 0 | 0.33 | **0.73** | **0.34** | **0.399** | 0.491 | -0.068 |
| 1v7m | 145 | 0 | 0 | 0 | 0.26 | 0.77 | 0.29 | **0.902** | **0.792** | **0.685** |
| 4g3y | 148 | **1** | 0.17 | 0.33 | 0.03 | 0.08 | 0.04 | 0.705 | **0.778** | **0.478** |
| 2vxt | 156 | 0.47 | 0.36 | 0.3 | 0.04 | 0.09 | 0.04 | **0.648** | **0.692** | **0.337** |
| 3u9p | 169 | 0.06 | 0.05 | 0 | 0.31 | **1** | **0.47** | **0.688** | 0.695 | 0.382 |
| 3o2d | 178 | 0 | 0 | 0 | 0.32 | 0.64 | 0.28 | **0.85** | **0.857** | **0.707** |
| 1fns | 196 | **1** | 0.07 | 0 | 0 | 0 | 0 | 0.839 | **0.727** | **0.555** |
| 3ma9 | 197 | 0 | 0 | 0 | 0 | 0 | 0 | **0.47** | **0.462** | -0.068 |
| 3rvv | 223 | 0.15 | 0.17 | 0.07 | 0.25 | **0.93** | **0.39** | **0.548** | 0.544 | 0.093 |
| 3raj | 230 | 0 | 0 | 0 | 0 | 0 | 0 | **0.519** | **0.521** | **0.04** |
| 1nfd | 239 | **0.92** | **0.7** | **0.75** | 0.07 | 0.23 | 0.04 | 0.565 | 0.639 | 0.19 |
| 3i50 | 273 | 0 | 0 | 0 | 0 | 0 | 0 | **0.893** | **0.819** | **0.708** |
| 3gjf | 276 | 0.05 | 0.11 | 0.05 | 0.15 | 0.66 | 0.2 | **0.64** | **0.759** | **0.381** |
| 3liz | 329 | 0 | 0 | 0 | 0.26 | **0.68** | **0.34** | **0.555** | 0.51 | 0.047 |
| 3pgf | 358 | 0 | 0 | 0 | 0.02 | 0.04 | 0 | **0.507** | **0.508** | **0.015** |
| 3zkm | 375 | 0 | 0 | 0 | 0.32 | **0.88** | **0.46** | **0.802** | 0.684 | **0.472** |
| 3r1g | 381 | 0 | 0 | 0 | 0.37 | **1** | **0.57** | **0.714** | 0.546 | 0.198 |
| 4jr9 | 409 | 0.46 | 0.5 | 0.46 | 0.19 | 0.85 | 0.34 | **0.657** | **0.901** | **0.502** |
| 4ene | 442 | 0 | 0 | 0 | 0 | 0 | 0 | **0.649** | **0.554** | **0.18** |
| 3o0r | 449 | 0 | 0 | 0 | 0.08 | **0.7** | **0.19** | **0.653** | **0.775** | **0.41** |
| 3t3p | 453 | 0.25 | 0.04 | 0 | 0 | 0 | 0 | **0.599** | **0.696** | **0.278** |
| 1n8z | 581 | 0 | 0 | 0 | 0 | 0 | 0 | **0.525** | **0.628** | **0.113** |

Table S1 The summarizing results on testing set.
